## Supplemental figures for "Phosphatidylserine-exposing medium/large extracellular vesicles: potential cancer biomarkers"

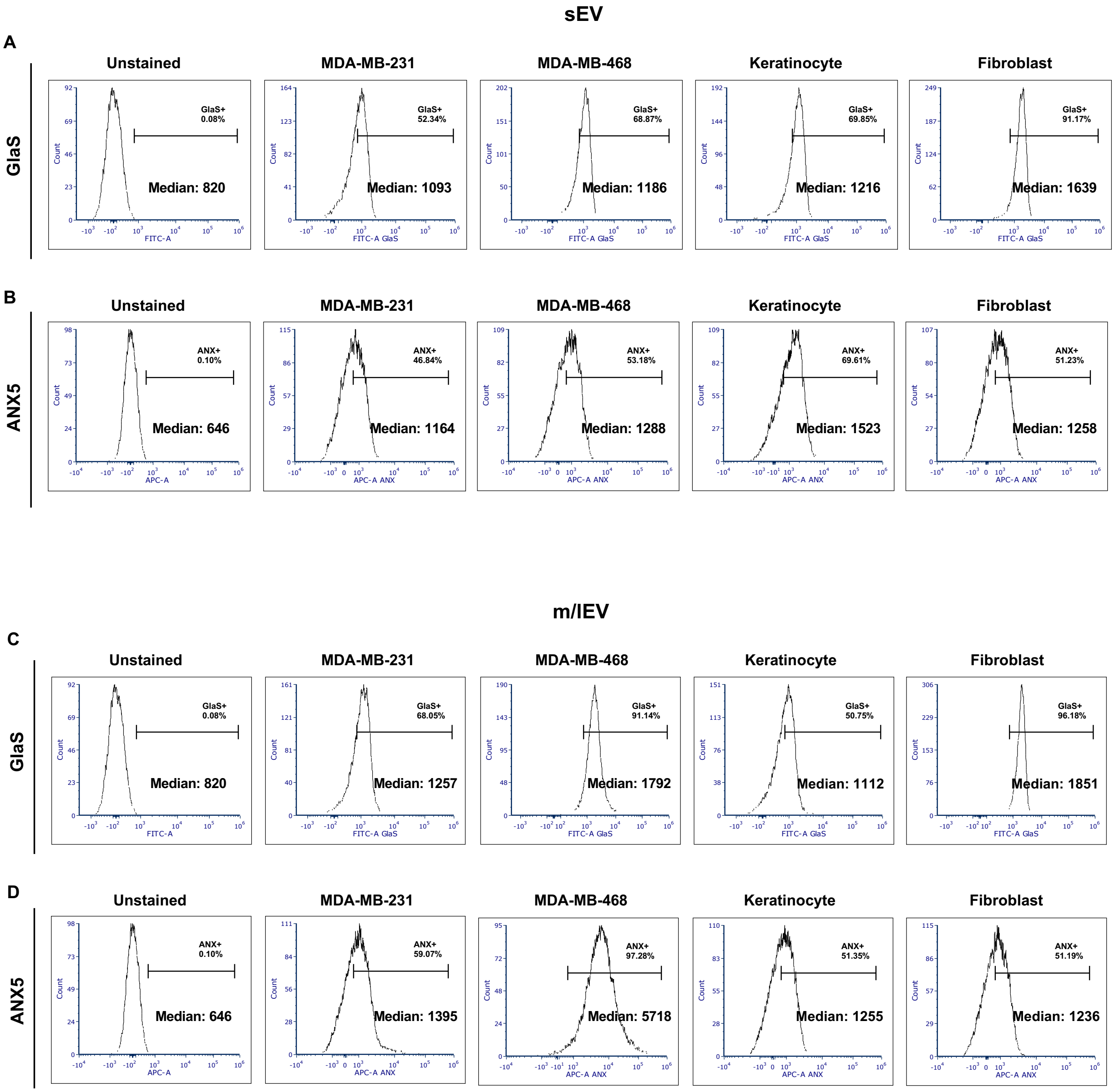


**Figure S1.** A representative of five analyses of PS exposure on the beads carrying sEVs or m/lEVs derived from cancer and non-cancerous cells. FITC-GlaS or APC-annexin A5 (ANX5) detected PS-exposing sEVs (A and B) or m/lEVs (C and D) on the beads. Marker gates denoting either % annexin A5+ (ANX+) or GlaS+ were set based on FMO control (unstained EV-loaded beads). Each histogram shows the median fluorescence intensity of FITC-GlaS or APC-ANX5 for singlet beads.

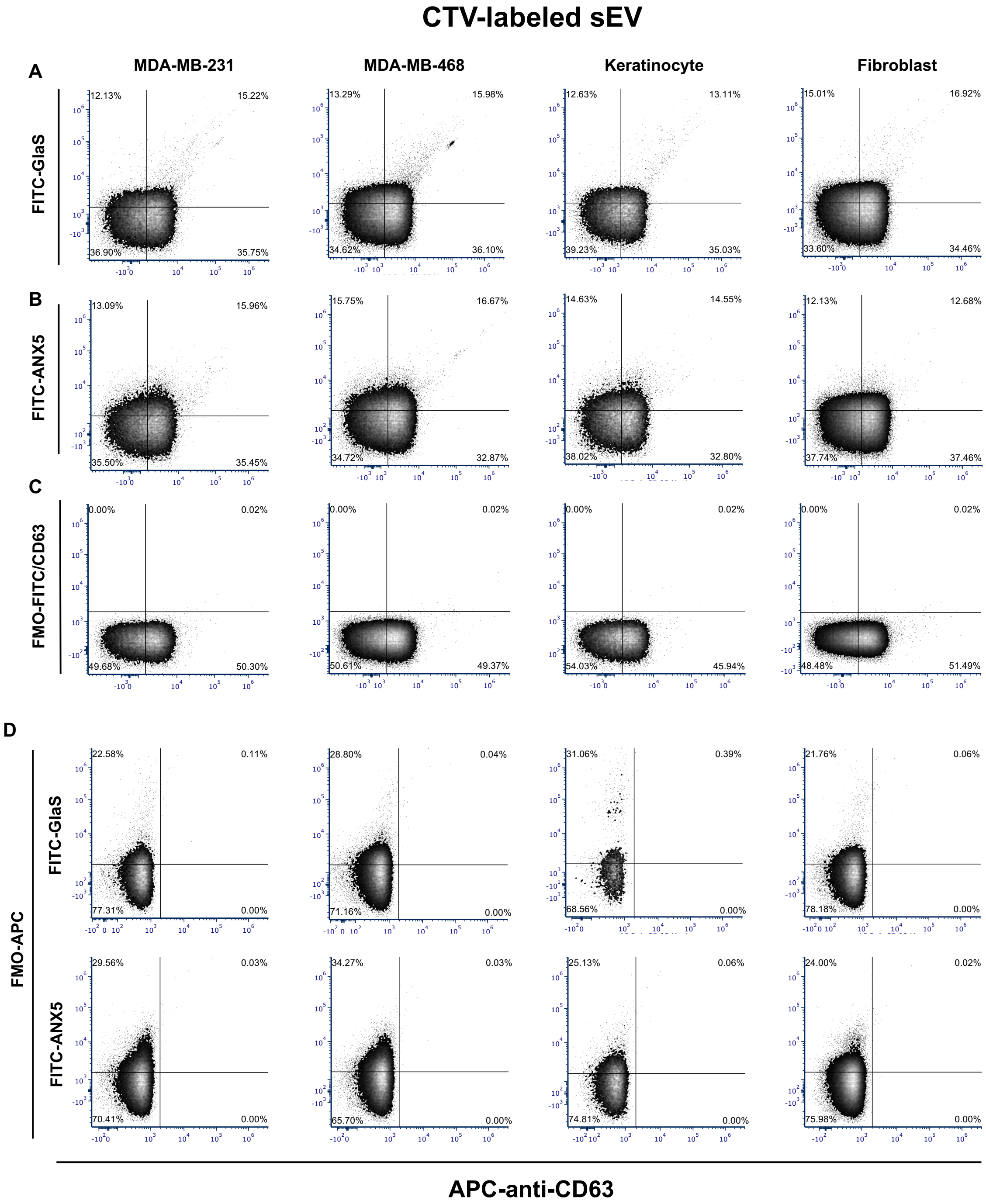

CTV-labeled sEV
MDA-MB-231
MDA-MB-468
Keratinocyte
Fibroblast
A
FITC-GlaS
12.13% 15.22% 36.90% 35.75%
13.29% 15.98% 34.62% 36.10%
12.63% 13.11% 39.23% 35.03%
15.01% 16.92% 33.60% 34.46%
B
FITC-ANX5
13.09% 15.96% 35.50% 35.45%
15.75% 16.67% 34.72% 32.87%
14.63% 14.55% 38.02% 32.80%
12.13% 12.68% 37.74% 37.46%
C
FMO-FITC/CD63
0.00% 0.02% 49.68% 50.30%
0.00% 0.02% 50.61% 49.37%
0.00% 0.02% 54.03% 45.94%
0.00% 0.02% 48.48% 51.49%
D
FMO-APC
FITC-GlaS
22.58% 0.11% 77.31% 0.00%
28.80% 0.04% 71.16% 0.00%
31.06% 0.39% 68.56% 0.00%
21.76% 0.06% 78.18% 0.00%
FITC-ANX5
29.56% 0.03% 70.41% 0.00%
34.27% 0.03% 65.70% 0.00%
25.13% 0.06% 74.81% 0.00%
24.00% 0.02% 75.98% 0.00%
APC-anti-CD63

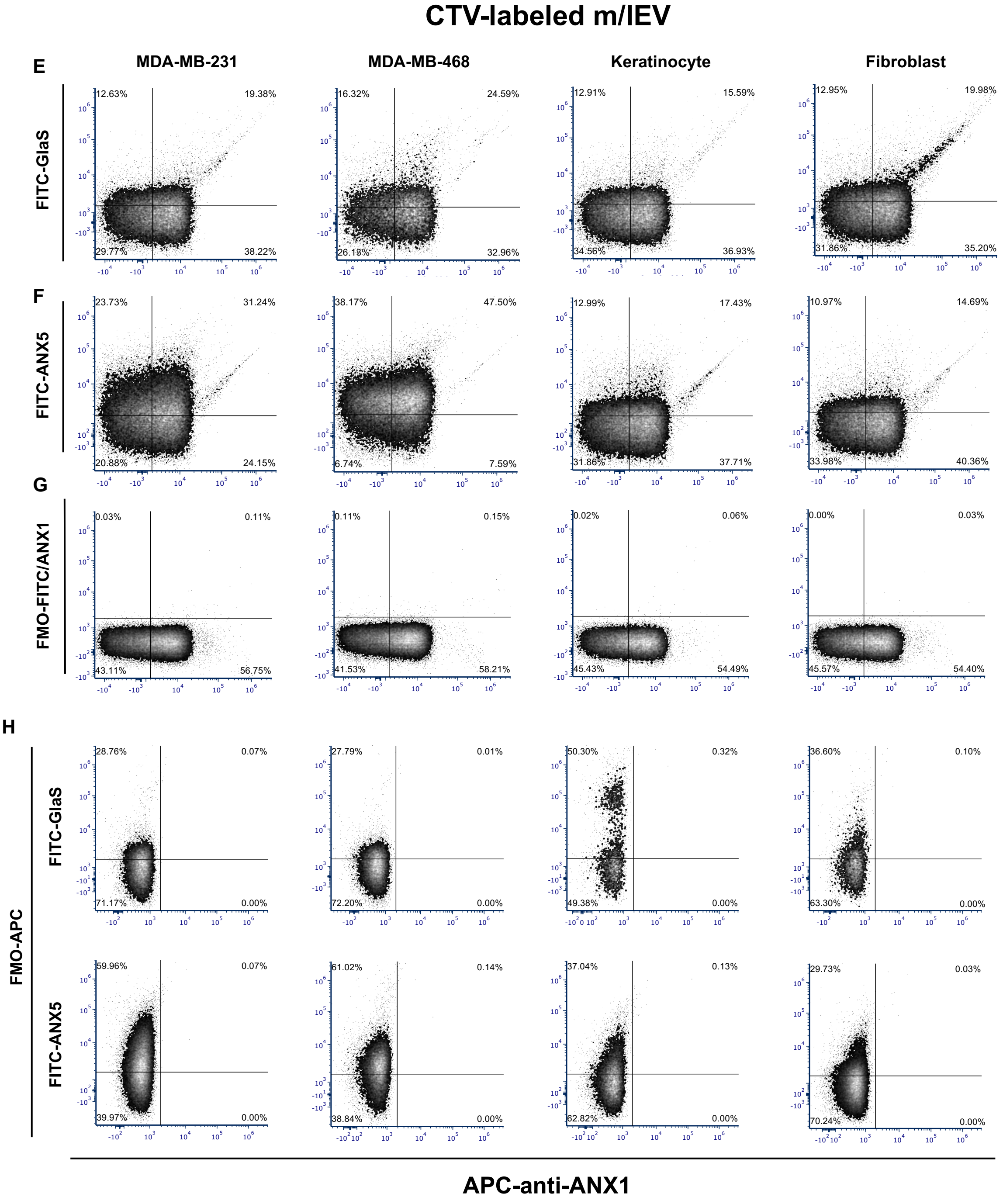


**Figure S2.** A representative of three independent analyses of PS-exposing sEVs and m/lEVs derived from cancer and non-cancerous cells. EVs were stained with CellTrace Violet (CTV). (A-D) CTV-sEVs were incubated with APC-anti-CD63 antibodies and FITC-GlaS or FITC-annexin A5 (ANX5). (E-H) CTV-m/lEVs were incubated with APC-anti-annexin A1 (ANX1) antibodies and FITC-GlaS or FITC-ANX5. FITC-annexin A5 (ANX5) or FITC-GlaS was evaluated to determine the frequency of individual PS-exposing EVs. Quadrant gates were set based on fluorescence minus one (FMO) controls (C, D, G, and H).
